# EvoFlow-RNA: Generating and Representing non-coding RNA with a Language Model

**DOI:** 10.1101/2025.02.25.639942

**Authors:** Sawan Patel, Fred Zhangzhi Peng, Keith Fraser, Adam D. Friedman, Pranam Chatterjee, Sherwood Yao

## Abstract

RNA plays a critical role across numerous biological functions. Recent advances in language modeling show promise with representing RNA, but the possibility of large-scale RNA design and optimization has not been fully explored. We propose **EvoFlow-RNA**, a bidirectional non-coding RNA language model leveraging a masked discrete diffusion model (MDM) formulation for both generative modeling and representation learning. EvoFlow-RNA bridges the gap between RNA sequence representation and design. It outperforms leading RNA models on three BEACON tasks critical to understanding RNA function, spanning from structure prediction to gene editing. For unconditional generation, it synthesizes diverse RNA sequences with native-like structural and binding properties. Additionally, EvoFlow-RNA can globally redesign aptamer sequences around preserved binding recognition sites with enhanced functionality. Our results demonstrate the effectiveness of EvoFlow-RNA in RNA modeling, highlighting the capability and potential of masked discrete diffusion for both recapitulating and enhancing existing RNAs.

## Introduction

Deep learning has revolutionized biomolecular design, serving as a foundational technology that drives innovation in biotechnology and medicine. Recent breakthroughs such as RFDiffusion [Watson et al., 2023], ProteinMPNN [Dauparas et al., 2022], and RosettaFold [Baek et al., 2021] exemplify the potential of computational approaches in protein representation with unprecedented *in silico* to *in vitro*, and *in vivo* success in modeling functional proteins. However, the majority of deep learning efforts are concentrated solely on protein understanding. By comparison, *de novo* RNA design —a field with immense application potential —has experienced less innovation. In addition to gene expression, RNAs play critical roles in gene regulation [Phizicky and Hopper, 2010, Olsen and Woese, 1993, Suzuki, 2021, Cai et al., 2009], catalysis [Dana et al., 2017], and information storage, making them key targets for many applications, such as RNA-based therapeutics, vaccine development, and programmable genetic circuits. The ability to design novel RNAs has opened new frontiers in therapeutic applications, including RNA-based drug delivery (e.g., gene therapies) and immune modulation [Coller and Ignatova, 2024, Hu et al., 2020, Kanasty et al., 2013, Morrow et al., 2024]. Synthetic RNA constructs, such as RNA aptamers, have shown significant potential in precision medicine, offering high-affinity and specific target binding for disease treatment and biomolecular recognition [Ng et al., 2006, Zhou et al., 2012]. As RNA-based technologies continue to evolve, their diverse functional repertoire underscores the need for advanced models tailored to enhancing RNA sequence and structure design.

Recent efforts in RNA modeling have begun integrating structural information to enhance generative and predictive capabilities. One such approach is a diffusion-based model that designs RNA aptamers with inherent fluorescent properties (i.e., via aptamer structure, not via chemical modification) and minimal sequence homology to known examples while maintaining functional integrity [Wong et al., 2024]. Despite their effectiveness, structure-based approaches are often constrained by the limited number of solved structures.

Sequence-based methods have also shown promise. BERT-style masked language models have gained traction for RNA representation learning, excelling in tasks such as functional annotation and mutation effect prediction [Penić et al., 2024, Chen et al., 2022, Zhang et al., 2024a, Chen et al., 2024, Akiyama and Sakakibara, 2022]. By pretraining on millions of non-coding RNA (ncRNA) sequences, these models learn meaningful embeddings that can be effectively transferred to downstream tasks. These include RNA-protein interaction prediction, structural stability assessment, and classification of regulatory elements. While powerful for prediction tasks, these models are not explicitly designed for sequence generation, limiting their utility in RNA design applications. By comparison, autoregressive language models have demonstrated several advantages in RNA design, achieving high fidelity in generating sequences with biologically meaningful properties [Zhang et al., 2024b, Zhao et al., 2024]. However, these models are inherently constrained by their left-to-right generation mechanism, limiting their ability to fully capture the bidirectional dependencies essential for RNA structure-function relationships. In particular, this deficit limits their ability in conditional generation tasks.

Masked discrete diffusion models, also known as masked diffusion models (MDMs), have emerged as a promising alternative to autoregressive models for the generative modeling of discrete data while also maintaining the bidirectional context formulation of BERT-like models. In MDMs, the forward process involves progressively masking elements of the input data, effectively adding noise by replacing original tokens with a mask token. The reverse process then aims to reconstruct the original data by predicting the masked tokens at each step, thereby learning the data distribution through iterative denoising. This approach has been shown to be effective in various applications, including language modeling and biological sequence generation [Shi et al., 2024, He et al., 2022, Wang et al., 2024, Peng et al., 2025]. In biological contexts, MDMs have comparable performance with leading BERT-based models in representation learning tasks [Hayes et al., 2025] in addition to improved generation quality compared to autoregressive language models [Peng et al., 2025, Sahoo et al., 2024, Manocha et al., 2021].

In this work, we propose EvoFlow-RNA (Figure 1), an RNA language modeling with bidirectional generative capabilities and representation learning. EvoFlow-RNA is an MDM, providing a unified framework for generation and representation by combining bidirectional attention and discrete flow matching. We evaluate EvoFlow-RNA in three key areas: representation learning, unconditional RNA generation, and RNA scaffold design.

**Fig. 1.**
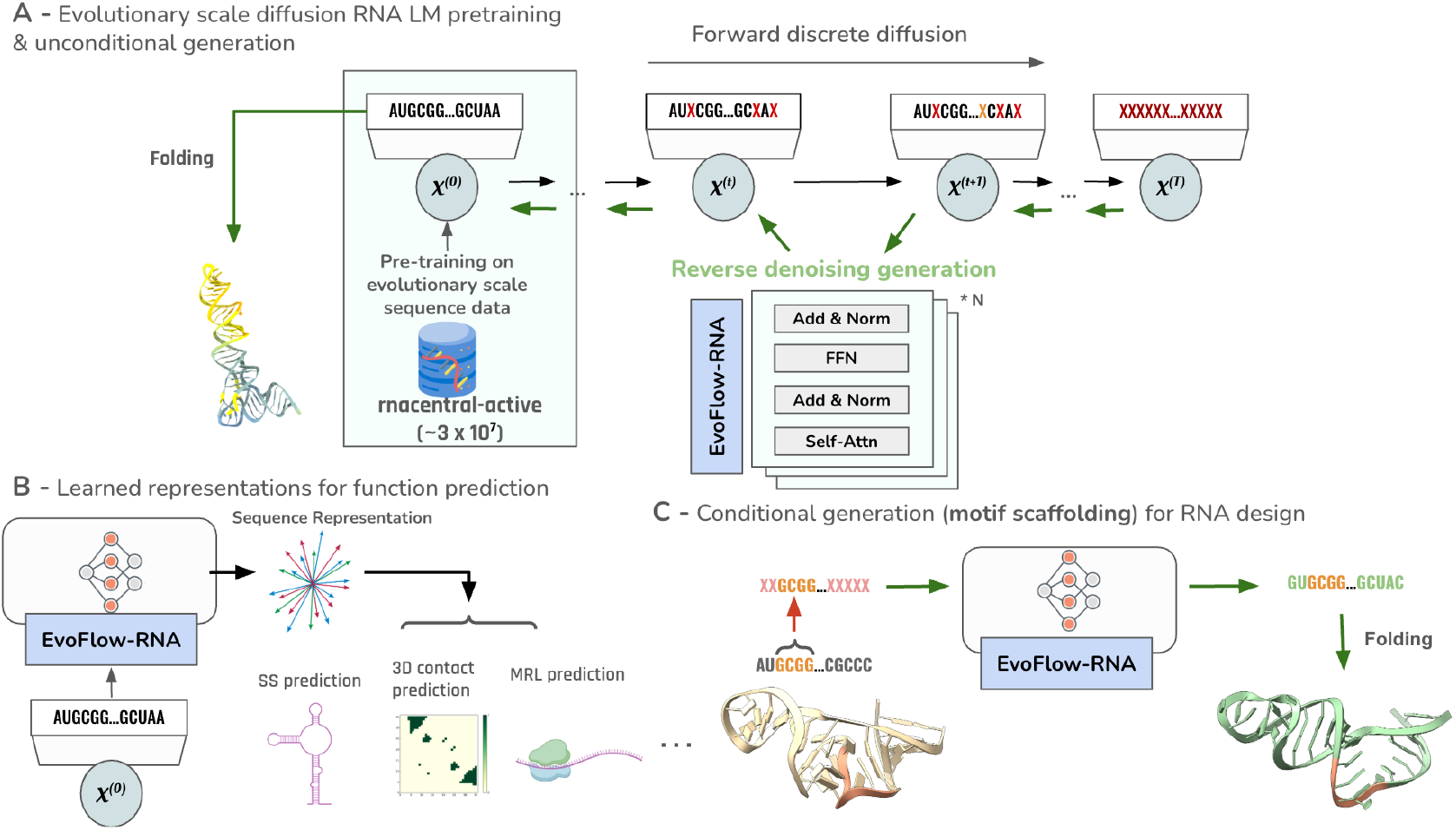
General framework of EvoFlow-RNA. A) Pre-training, modeling and unconditional generation. B) Non-coding RNA representation learning for structure, functional and engineering design tasks. C) Conditional generation of RNAs via motif scaffolding.

In representation learning, EvoFlow-RNA is benchmarked on 13 downstream tasks from the BEACON suite [Ren et al., 2024]. Across the board, our model matches and, in some cases, outperforms baseline methods. It excels in structural prediction, particularly secondary structure and contact map prediction, and on a variety of RNA engineering tasks. This demonstrates the utility of masked diffusion latent representations.

For unconditional RNA generation, EvoFlow-RNA generates *de novo* ncRNA sequences with biophysical properties closely aligned to native sequences, outperforming an autoregressive counterpart and other relevant baselines. Across various lengths, it maintains consistency in key metrics such as pLDDT, GC content, minimum free energy, and sequence entropy. Notably, it inherently reproduces common folds found in well-studied RNA classes, such as tRNAs.

In RNA scaffolding, EvoFlow-RNA’s bidirectional generation conditionally designs aptamer sequences while preserving key binding motifs. It successfully redesigns aptamers binding to HIV-1 Rev peptide and AMP, maintaining functional elements with motif-specific RMSDs below 1Å while exhibiting novel global conformations. Moreover, these redesigned aptamers often exhibit higher docking scores than their wild-type counterparts. This is indicative of heightened binding capability. As follows, we show that EvoFlow-RNA maintains well-formed representations of ncRNAs in its latent space while having an unparalleled ability for *de novo* generation.

## Methods

### Background

In this section, we describe the key components of the Discrete Flow Model (DFM) that underlies EvoFlow-RNA. We first introduce the notation and problem formulation, then describe the construction of the generative flow using continuous-time Markov chains (CTMCs). We detail our method formulation with special emphasis on the mask formulation, and finally explain the sampling procedure and training objective.

### Notation

Let *x* ∈ {1, …, *S*}^*D*^ denote a discrete sequence with *D* dimensions, where each element takes one of *S* possible states. For ease of exposition, we assume *D* = 1; the extension to higher dimensions follows by appropriate factorization.

We define *p*_0_(*x*) as the initial (noise) distribution, *p*_data_(*x*) as the target data distribution, and *p*_*t*_(*x*) as the time-dependent marginal distribution for *t* ∈ [0, 1] that interpolates between *p*_0_ at *t* = 0 and *p*_data_ at *t* = 1. The dynamics of *p*_*t*_(*x*) are generated by a time-dependent rate matrix *R*_*t*_ ∈ R^*S*×*S*^, whose off-diagonal entries are non-negative. In the context of continuous-time Markov chains (CTMCs), the transition probability for an infinitesimal time step Δ*t* is given by

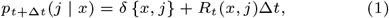

where *δ* {*x, j*} is the Kronecker delta, and *R*_*t*_(*x, x*) is defined so that the row sums are equal to 1. We also introduce a conditional noise flow *p*_*t*|1_(*x*_*t*_ | *x*_1_) that interpolates from a prescribed noise distribution *p*_noise_(*x*) at *t* = 0 to a data point *x*_1_ at *t* = 1.

#### Problem Formulation

Our objective is to design a generative model that transforms noise samples into samples from the data distribution. More precisely, we seek a *generative flow* {*p*_*t*_(*x*)}_*t*∈[0,1]_ satisfying

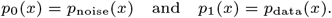

*p*_0_(*x*) = *p*_noise_(*x*) and *p*_1_(*x*) = *p*_data_(*x*).

The evolution of the marginal *p*_*t*_(*x*) is governed by the Kolmogorov forward (or continuity) equation:

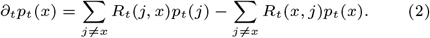

In our approach, the generative flow is constructed via an expectation over conditional flows:

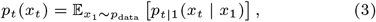

which decomposes the global evolution into simpler, datapoint-conditional processes.

#### Method Formulation

A key component of our approach is the explicit construction of the conditional flow *p*_*t*|1_(*x*_*t*_ | *x*_1_). We consider two variants: a uniform corruption flow and a mask-based flow. In this work, we focus on the mask formulation, which has been found effective for discrete data.

Mask Formulation The mask formulation defines the conditional flow as

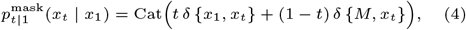

where *M* is an artificially introduced mask state. This construction satisfies the following limits: at t = 0, 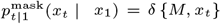, so that all probability mass is on the mask state, and at *t* = 1, 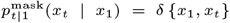, thereby recovering the true data point. Using (4), the overall generative flow in (3) becomes

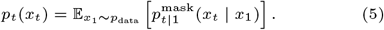

In order to sample from *p*_*t*_(*x*_*t*_), we require a rate matrix *R*_*t*_(*x*_*t*_, *j*) that generates this flow. One strategy is to first derive a conditional rate matrix *R*_*t*_(*x*_*t*_, *j* | *x*_1_) that produces the desired *p*_*t*|1_(*x*_*t*_ | *x*_1_) and then aggregate it via the denoising distribution:

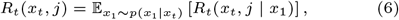

With

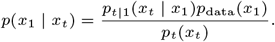

This formulation decouples the design of the conditional dynamics (which can be chosen in closed form) from the unconditional generative process.

*Sampling*

To sample from the data distribution, we simulate the CTMC defined by the rate matrix *R*_*t*_(*x*_*t*_, *j*). Starting from an initial sample *x*_0_ ∼ *p*_0_(*x*), we discretize the time interval [0, 1] with a step size Δ*t* and update the state according to

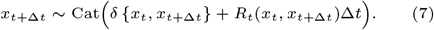

This process is repeated iteratively until *t* = 1, at which point the final state is a sample from *p*_data_(*x*).

#### Training

The parameters of our model are learned by training a neural network 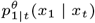 to approximate the denoising distribution *p*(*x*_1_ | *x*_*t*_). The training objective is a standard cross-entropy loss:

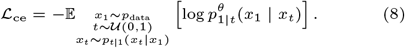

Here, *t* is sampled uniformly from [0, 1] and *x*_*t*_ is generated using the conditional flow (e.g., via the mask formulation in (4)). Importantly, this loss does not depend on the choice of the conditional rate matrix *R*_*t*_(*x*_*t*_, *j* | *x*_1_), which allows flexibility in designing the sampling dynamics.

### Dataset

RNAcentral is a comprehensive database that consolidates ncRNA sequences from over 40 expert databases, providing a unified platform for ncRNA data. As of its latest release, RNAcentral encompasses more than 30 million ncRNA sequences spanning a diverse range of organisms and RNA types. We take our pretraining data from RNACentral [Consortium, 2021], which has a processed dataset of over 30 million ncRNAs. We made no further preprocessing adjustments.

### Tokenization

Each nucleotide is treated as a single token, following a character-level tokenization scheme specifically designed for RNA sequences. During tokenization, all uracil (U) bases are replaced with thymine (T) to maintain consistency across sequence representations, resulting in a primary vocabulary of four nucleotides: “A”, “C”, “T”, and “G”. Additionally, we include IUPAC degenerate nucleotide codes such as “I”, “R”, “Y”, “K”, “M”, “S”, “W”, “B”, “D”, “H”, “V”, “N”, and “-” to account for sequence ambiguities commonly found in biological datasets. To align with standard masked language modeling (MLM) practices, we incorporate the special tokens [CLS], [EOS], [MASK], and [PAD], which respectively denote sequence classification, end-of-sequence markers, masked tokens for pretraining, and padding for fixed-length sequences. EvoFlow-RNA processes sequences of up to 1022 tokens, dynamically truncating or padding them as needed. Random sequence cropping is applied during training to ensure the model observes diverse sequence contexts across epochs, which enhances its ability to generalize across RNA families.

### Encoder Architecture

EvoFlow-RNA is a BERT-style encoder-only transformer that extends the RiNALMo-150M model [Penić et al., 2024] and is trained on a masked diffusion modeling objective while leveraging the same tokenization scheme and architectural framework. Given an input sequence *x* = (*x*_1_, *x*_2_, …, *x*_*L*_) of length *L*, the model maps it to a sequence of context-aware token embeddings of shape (*L*×*M*_emb_). The architecture consists of several transformer blocks, where each block contains a multi-head self-attention (MHSA) module with 20 attention heads, a feed-forward network (FFN) using the SwiGLU activation function [Shazeer, 2020], and Layer Normalization (LayerNorm) within residual connections to stabilize training dynamics. EvoFlow-RNA adopts Rotary Positional Embeddings (RoPE) [Su et al., 2024] to encode both absolute and relative positional information, thereby improving its ability to model long-range dependencies in RNA sequences. To enhance computational efficiency, we employ FlashAttention-2 [Dao, 2024], a memory-optimized exact attention mechanism that significantly reduces peak activation memory without compromising precision, enabling training on long RNA sequences with high batch throughput.

EvoFlow-RNA is initialized from all three available pretrained RiNALMo checkpoints (33M, 150M, 650M) and continues training using a masked diffusion objective with a token corruption strategy where *t* ∼ U (0, 1) of the input sequence is masked, namely, replaced by [MASK]. This approach ensures that the model retains the strong representation learning capacity of RiNALMo while being effectively adapted for diffusion-based sequence generation. Compared to GELU-based architectures, the incorporation of SwiGLU activation improves gradient stability, while RoPE enables better positional encoding compared to conventional absolute position embeddings. Notably, we observe significant memory savings with FlashAttention-2 while maintaining model convergence, making EvoFlow-RNA computationally efficient for large-scale RNA sequence modeling. Configurations can be found in Table A1.

### Diffusion Training

We take two-stage training as described in Manocha et al. [2021], which involves the initial training of a masked language model (MLM) followed by continuing that training with a masked diffusion modeling (MDM) objective. We truncate all RNAs to 1022 tokens and have no further data processing. Our model is trained for 100K updates, with the batch size set at 320K for the 33M and 150M parameter models and 1M for the 650M parameter model. We additionally utilize a batch sampler to construct batches using sequences of similar lengths. Practically, we did not implement the first-stage training. Instead, we employed RiNALMo, an RNA language model that is trained on MLM loss. We finetune RiNALMo using the MDM objective with the weighted cross-entropy loss where the weight is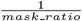, and the *mask ratio* is uniformly sampled from 0 to 1 and clamped to 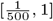.

### Sampling

We follow the P2 sampling scheme from Peng et al. [2025], specifically self-planning. In P2, the denoiser makes predictions and the planner selects positions to update (unmask to the prediction or remask an existing token). In self-planning, the planner is the denoiser itself where we leverage the denoiser’s predicted probabilities to score positions and unmask the mask tokens that have the top predicted probabilities and remask the unmasked tokens that have the lowest. The overall sampling scheme is depicted in algorithm 1.

**Algorithm 1.**
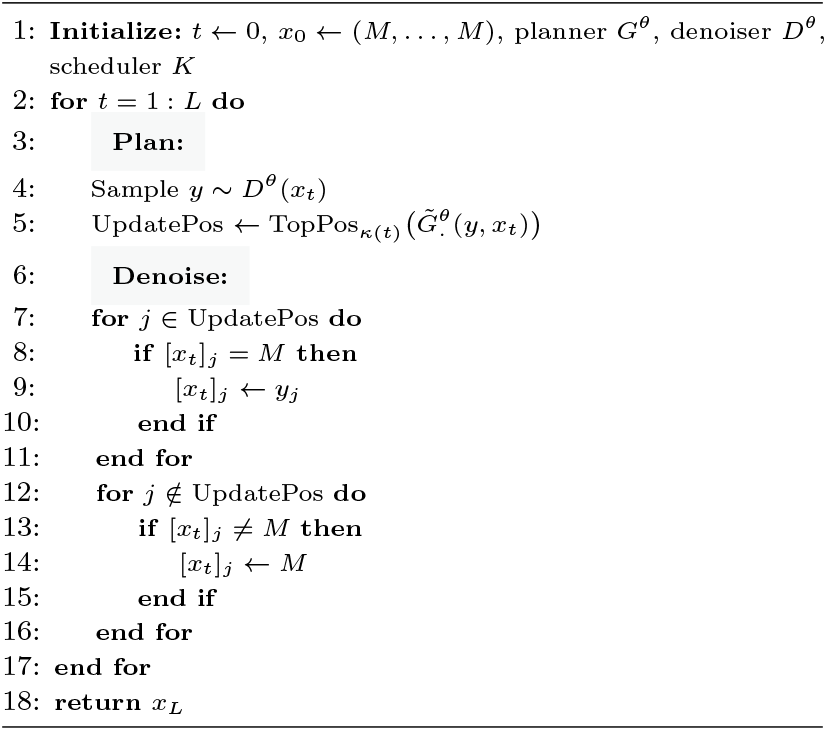
Sampling Scheme.

### In silico validation

An evaluation of the generative capability of the EvoFlow RNA model was performed by comparing the binding capabilities of 6 designer RNA aptamers to wild type variants. Here we used three aptamers that were able to bind to fluoroquinolone derivatives (PDB ID: 7FHI), AMP (PDB ID: 1RAW), and HIV inhibitor (PDB ID: 2L94). We also used three aptamers that bind to HIV-1 peptide (PDB IDs: 1ULL, 1ETG), and the RSG peptide (PDB ID: 1I9F). To evaluate binding of both wild type and designer aptamers to these targets we used HADDOCK version 3.0 [M.C. Teixeira et al., 2024]. Structures were retrieved from the PDB, and missing residues were analyzed and refined using Swiss-PdbViewer version 4.1.0. The simulations generated multiple binding poses across several clusters, each with an associated docking score that is representative of the strength of the binding interaction between the ligand (small molecule, and peptide) and the RNA aptamers. In this study we used the calculated binding energy of the wild type structures as the baseline to compare against out designed candidates. Poses and binding energies calculated for cluster 1 of each simulation were used for this comparison as they represented conformations of the ligand in the wild-type and generated aptamers in similar binding pockets.

### Evaluation Metrics

Across both representation learning and unconditional generation, we utilize a wide array of metrics for assessing the performance of EvoFlow-RNA. Metrics used in BEACON representation learning are defined as the yare in BEACON [Ren et al., 2024]. pLDDT is the predicted local distance difference test, a per-residue measure of local confidence from 0 to 100 (or normalized to span from 0 to 1) [Tunyasuvunakool et al., 2021]. pLDDT values are computed via RhoFold [Shen et al., 2024]. Higher scores indicate a more confident and, typically, accurate prediction. GC content is the percentage of guanine (G) or cytosine (C) nucleotides within a nucleic acid sequence. MFE, or minimum free energy, is the predicted minimum free energy of an RNA which indicates the most stable state of a system at equilibrium. MFE computations are taken from ViennaRNA [Lorenz et al., 2011]. Sequence entropy is the Shannon entropy of a sequence, a measure of predictability or, consequently, diversity [Shannon, 1948].

## Results

### Representation Benchmarking

We evaluate EvoFlow-RNA representations using the BEACON benchmark suite [Ren et al., 2024]. Accordingly, the model is fine-tuned and assessed for performance across 13 related tasks spanning three categories: structure, function and engineering. We compare our model’s performance against leading pretrained large-language models (LLMs) and supervised models. The supervised models include a convolutional neural network (CNN), ResNET, and LSTM. The pretrained LLMs include RNA-FM [Chen et al., 2022], RNA-BERT [Akiyama and Sakakibara, 2022], RNA-MSM [Zhang et al., 2024a], SpliceBERT [Chen et al., 2024], 3UTRBert [Yang et al., 2024], UTR-LM [Chu et al., 2024], BEACON-B [Ren et al., 2024], RNAGenesis, and [Zhang et al., 2024b]. We also include results for RiNALMo [Penić et al., 2024] for direct comparison to EvoFlow-RNA. These models vary widely in size, up to 27M parameters for the supervised models and up to 866M parameters for the pretrained LLMs. They are also pretrained on different data sources, including but not limited to ncRNA, pre-mRNA, mRNA-3’UTR, and mRNA-5’UTR. For EvoFlow-RNA, we utilize the exact configurations listed in [Ren et al., 2024] for each task to ensure consistency. The only exceptions are secondary structure prediction, contact map prediction and distance map prediction, where we were not able to train with a batch size of 32 due to hardware limitations. We instead trained with a batch size of 2, with all other hyperparameters held constant.

In Table 1, we show the performance of EvoFlow-RNA against all models evaluated against the BEACON benchmark, pulled across BEACON [Ren et al., 2024] and RNAGenesis [Zhang et al., 2024b]. Evidently, we observe that EvoFlow-RNA achieves state-of-the-art performance on 3 of the 13 evaluation tasks, reflecting the quality of masked diffusion model representations in the context of ncRNA. Interestingly, EvoFlow-RNA’s benchmarking scores nearly parallel those of its RiNALMo predecessor, showing that masked diffusion fine-tuning does not result in representation learning performance deterioration. Additionally, these models together appear to outperform RNAGenesis across several tasks, a model nearly 6 times as large.

**Table 1.**
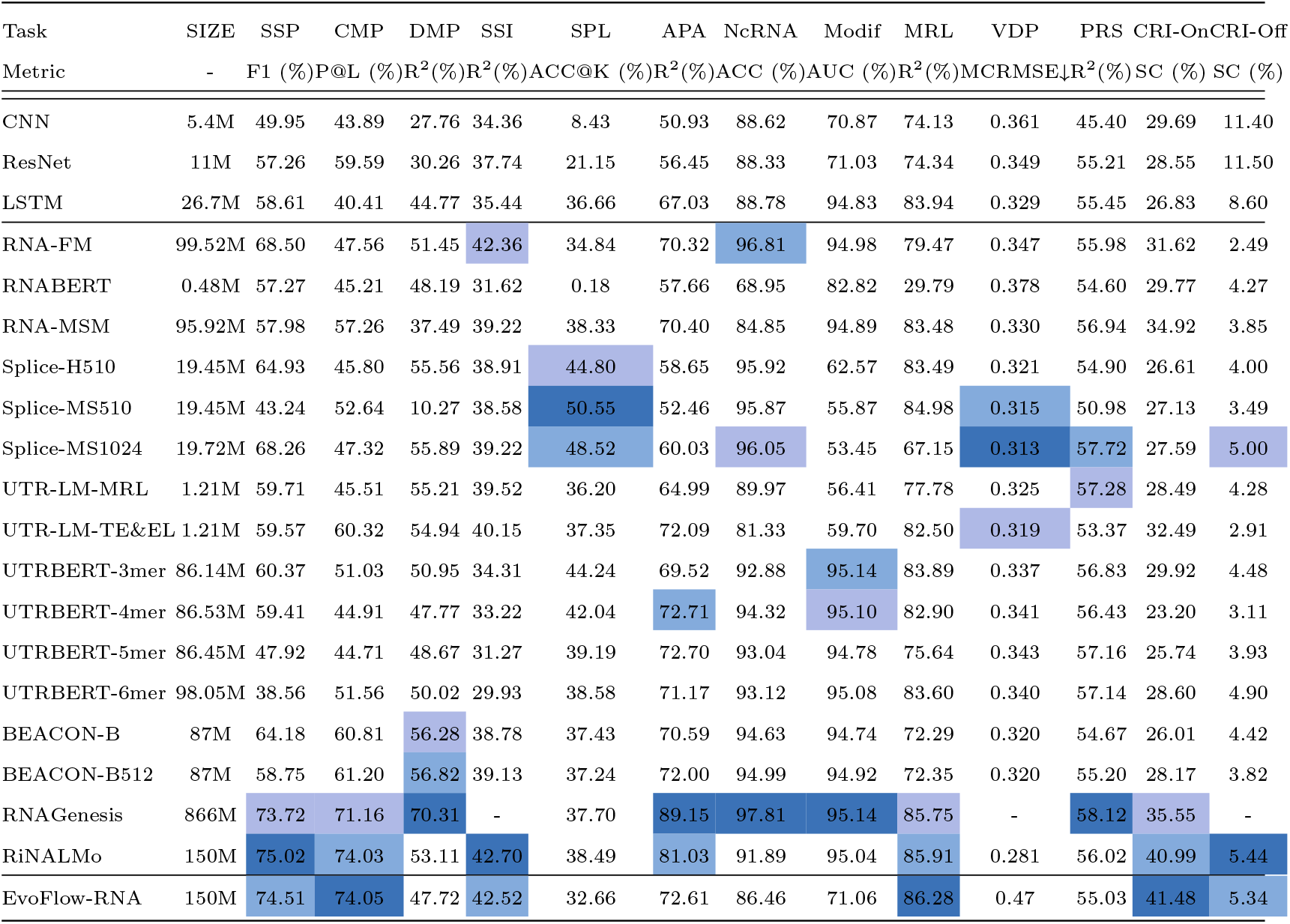
BEACON Representation Learning. Benchmarking across 13 downstream RNA-related tasks across structure, function and engineering. EvoFlow-RNA is compared with both naive supervised methods and pretrained RNA language models.

Notably, EvoFlow-RNA performs well at the structure-related tasks, such as secondary-structure prediction (SSP) and contact map prediction (CMP). The latter requires an understanding of the three-dimensional representation of an RNA, showing that this property can be learned to a degree from sequence. Interestingly, though the model performs well at contact map prediction, it does not perform as well at distance map prediction (DMP), another task requiring a precise understanding of RNA tertiary structure. Though, this drop-off is not a significant departure from the general trend exhibited by all other models. Regardless, we maintain that EvoFlow-RNA retains the rich latent representations inherited from its pretrained BERT checkpoint, in addition to its knack for assessing the functional capabilities of ncRNAs such as single-guide RNAS for CRISPR on-target gene editing (CRI-On) and predicting the mean ribosomal load of a given mRNA sequence (MRL).

Lastly, we also show that model performance on secondary structure prediction scales with complexity as in Table A2.

### ncRNA de novo Generation Evaluation

#### Unconditional generation

Synthetic ncRNAs - viable tools in therapeutics, diagnostics and biotechnology - can encompass engineered tRNAs, codon-optimized mRNAs, or siRNA therapeutics. Moreover, RNA aptamers specifically have a long history of utility as short RNA molecules that can bind to specific biomolecules with high affinity, selectivity, and specificity. We evaluated EvoFlow-RNA’s ability to design ncRNAs that align with key properties computed from the ncRNA training dataset. This would allow for the generation of a diverse library of potential RNA aptamers, each exhibiting unique structure-stability profiles. These libraries would also diminish the need for the multiple rounds of SELEX required by canonical aptamer design.

To evaluate the model’s generative capability, we compared EvoFlow-RNA with GenerRNA [Zhao et al., 2024], a 350M parameter autoregressive model, and two RiNALMo checkpoints (150M and 650M). We generated aptamers from RiNALMo using RDM sampling. Lastly, we also featured native ncRNAs as a baseline for comparing key properties of the generated sequences. Native ncRNAs refer to ncRNAs found in living organisms, which were randomly drawn from the training dataset. To facilitate comparisons, we computed pLDDT as a proxy for foldability, GC content, minimum free energy, and sequence entropy. Libraries of size 50, 100, 200 and 400 were generated to faciliate this study. 100 sequences are generated per library size and per model. Per Figure 2, we report that EvoFlow-RNA designed ncRNAs exhibit similar biophysical profiles as native sequences than either of the RiNALMo base models. This is evident across all metrics, most notably GC content, MFE, and sequence entropy. In comparison to GenerRNA, EvoFlow-RNA produces more ‘foldable’ ncRNAs, as exhibited by the higher pLDDT distribution at the same level of sequence diversity. The EvoFlow-RNA pLDDT distribution is also relatively higher than the native sequence library, which may be a byproduct of oversampling particular sequences with higher foldability. More extensive results across library sizes are shown in Figure A1.

**Fig. 2.**
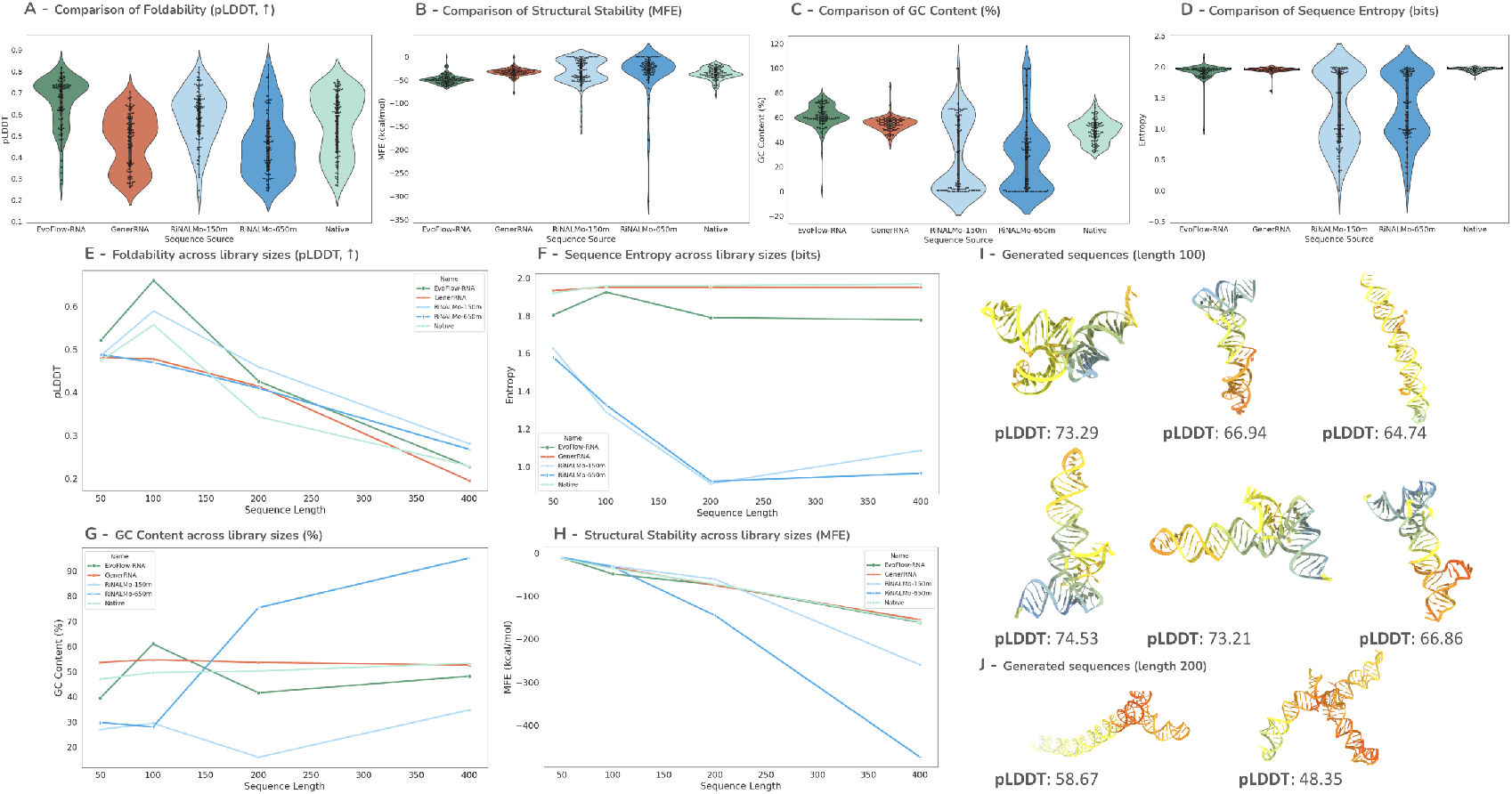
Unconditional generation evaluation with P2 (RDM, *η* = 1.0). Distributions for pLDDT, predicted minimum free energy, GC content, and sequence entropy for libraries generated from five sources. Featured are EvoFlow-RNA, a masked diffusion model, GenerRNA, an autoregressive model, two models of varying size for RiNALMo, a BERT model, and native ncRNA sequences from nature. (A-D) Metrics for sequences of length 100. E) Average pLDDT, F) MFE, G) GC content, and H) sequence entropy computed for libraries of size 100 as a function of library sequence length. De novo, unconditionally-generated sequences of I) length 100 and J) 200. Bases are colored according to their pLDDT and the AlphaFold2 color scheme. Additional library analysis and predicted structures can be found in Figures A1 and A2, respectively.

Most notably, we show that the consistency in alignment between EvoFlow-RNA and native ncRNAs holds across all evaluated sequence lengths, regardless of metric. All models converge to low average pLDDT values as the sequence length increases, which is likely a product of the folding model. *De novo* designs also exhibit a variety of tertiary structures, with several exhibiting tRNA-like structures per Figures 2 I,J and A2.

To assess sampling quality, we performed an ablation using various P2-derived strategies using three generation sources: EvoFlow-RNA (denoted as MDM), RiNALMo-150M and RiNALMo-650M in Table 2. We compared four sampling cases: ancestral (P2, *η* = 0), RDM (P2, *η* = 1), P2 with a calibrated *η*, and P2 with a dedicated BERT planner (RiNALMo-150M). It is apparent that training an MDM overall yields higher quality *de novo* ncRNAs over a pretrained BERT model such as RiNALMo, regardless of model size. Notably, ncRNAs generated from EvoFlow-RNA are comparable to a set of subsampled native ncRNA sequences, as per the objective metrics. The only exception, as also shown above, is pLDDT where sequences generated synthetically tend to exhibit higher values.

**Table 2.** RNA Sequence Generation Benchmark. The “Native” row represents subsampled natural RNA sequences. “MDM” refers to a pretrained 150M Masked Diffusion Model trained on RNACentral [Consortium, 2021].

| Sequence Source | pLDDT (↑) | MFE (kcal/mol) | Entropy | GC Content (%) |
| --- | --- | --- | --- | --- |
| Native | 48.26 | -35.83 | **1.96** | 49.64 |
| RiNALMo-150M | 59.01 | -30.12 | 1.29 | 29.50 |
| RiNALMo-650M | 46.99 | -31.90 | 1.33 | 28.06 |
| EvoFlow-RNA + Ancestral | 68.12 | -48.46 | 1.93 | 60.84 |
| EvoFlow-RNA + RDM | 67.35 | -47.54 | 1.89 | 59.42 |
| **EvoFlow-RNA + P2 (self-planning)** | **69.41** | -48.21 | 1.89 | 59.84 |
| **EvoFlow-RNA + P2 + RiNALMo-150M Planner** | **73.28** | **-51.91** | 1.86 | **65.47** |

Moreover, we also demonstrate that P2 sampling with a supervisory planner yields sequences with ‘best’ performance across all metrics, though we note the additional deviation from native sequences in the training dataset.

#### Conditional generation

A noteworthy ability of masked diffusion models, as opposed to their autoregressive counterparts, is the ability to inform generation using bidirectional context. This property is, in theory, useful for building scaffolds around key binding motifs. In this task, we fix one or more key motif subsequences within an experimentally validated aptamer with a crystal structure as a prompt and use EvoFlow-RNA to build around it. This task is particularly suited for masked diffusion models because bases upstream of the fixed motif can be generated in the context of the fixed motif.

To test this capability, we investigated two known RNA aptamers from the protein data bank: an RNA aptamer from Ye et al. [1996] that binds selectively to the HIV-1 Rev peptide (1ULL, length 35), and an RNA aptamer from Dieckmann et al. [1996] that binds selectively to the AMP molecule (1RAW, length 36). Within their respective depositions, we identified the key recognition sites of each aptamer that participate in the binding complex with the intended target. The subsequences corresponding to these recognition sites were fixed (bases 3:12, 21:23, 25:28 and 30:33 for 1ULL and bases 7-11 for 1RAW). The remainder of the sequence was masked and infilled using EvoFlow-RNA. Generated aptamers from these scaffolds were of the same length as the original.

Using this procedure, we designed a library of model-scaffolded aptamers. The result is shown in Figure 3, where we feature two conditionally-generated aptamers for each original aptamer. All scaffolded aptamers maintain similar MFE and GC content profiles as the original aptamer. Notably, the scaffolded aptamers for HIV-1 Rev peptide align fairly well with the original aptamers, with motif-specific RMSDs of approximately 1 Å and whole-structure RMSDs of approximately 4 Å, following a least-squares fit. The common motif RMSDs for the scaffolded and original AMP aptamers are below 1 Å, highlighting that the key recognition elements do not deviate substantially from their original locations using a scaffolding approach. However, we do note that the whole-structure RMSDs following a least-squares fit are well above 10 Å. This indicates that the conditionally-generated aptamers have a globally unique structure in comparison to the originals, though they maintain similarities at the local-level, and particularly at binding sites. Additional scaffolds for the HIV-1 Rev peptide system can be found in Figure A3.

**Fig. 3.**
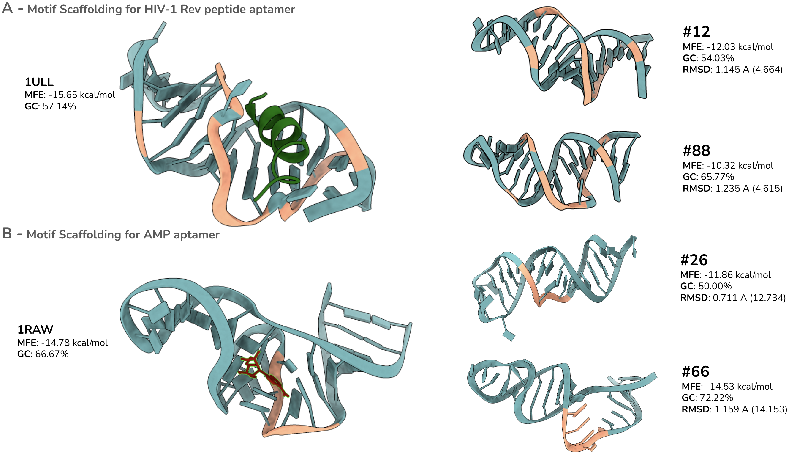
Conditionally generating scaffolds for RNA aptamers. A) 1ULL (Left), an RNA aptamer/HIV-1 rev peptide complex, is taken as a simpler study with several bases contributing to the binding recognition site (salmon) on the aptamer. (Right) Two conditionally-generated scaffold RNA aptamers sharing the binding recognition site (salmon) of the original aptamer. B) 1RAW (Left), an RNA aptamer/AMP molecule complex, is a more complicated study with fewer bases contributing to the binding recognition site (salmon) on the aptamer. (Right) Two conditionally-generated scaffold RNA aptamers sharing the binding recognition site (salmon) of the original aptamer.

#### Docking

Molecular docking is a valuable tool for the rapid evaluation of protein-ligand binding interactions as it provides us with the ability to compare binding poses and elucidate the molecular fingerprints at binding interfaces. In this study, we demonstrate that docking with HADDOCK version 3.0 enabled us to recapitulate binding of the target to the RNA aptamer in the same binding pocket observed in the crystal structure, albeit with slightly different conformations in some instances. We used associated binding energy (kcal/mol) to compare our generated aptamers to wild-type analogs in Figure 4.

**Fig. 4.**
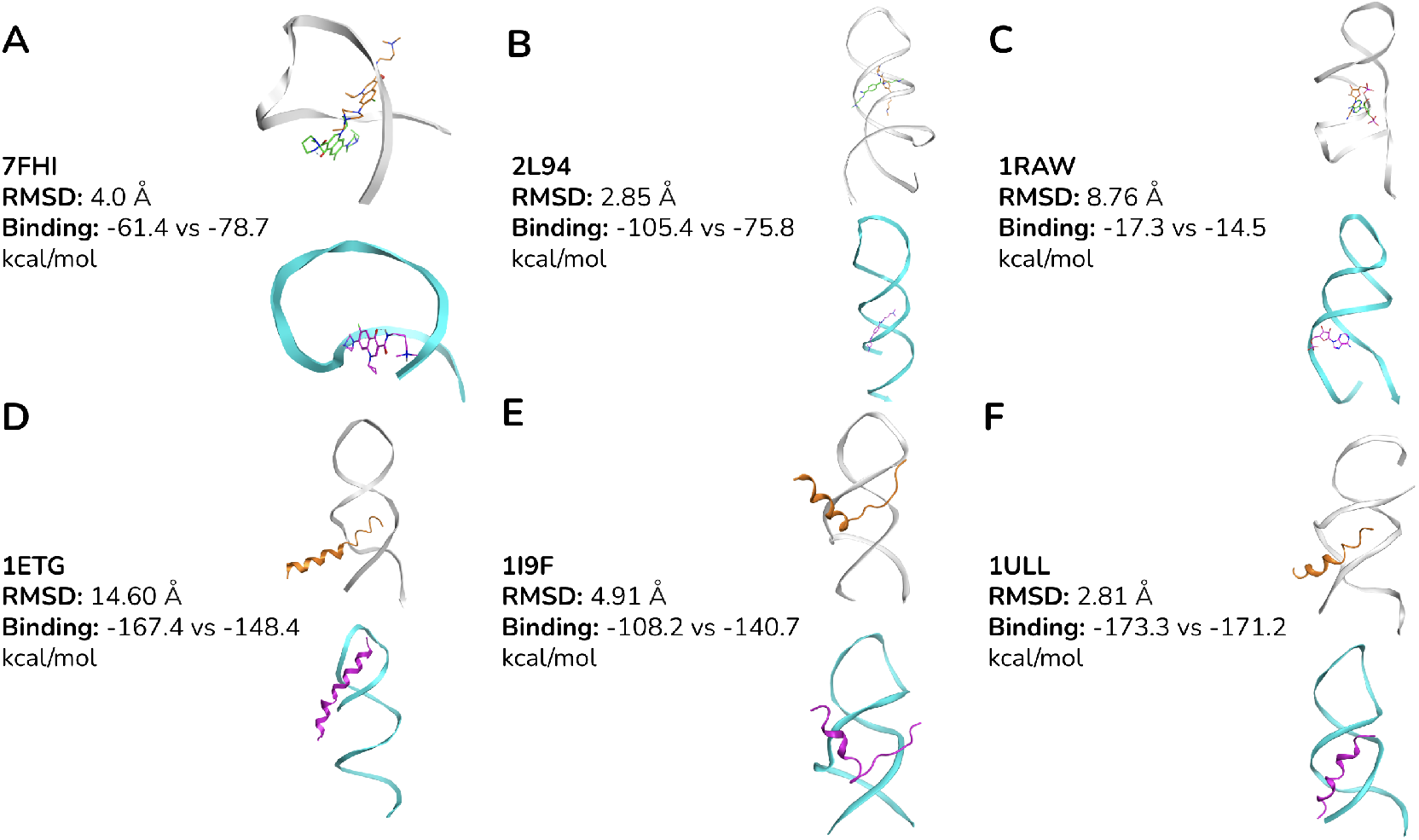
Haddock 3.0 binding scores. WT aptamers are shown in gray and model-scaffolded mutants are in cyan. Docking results are shown for three aptamer-small molecule systems (7FHI, 2L94, 1RAW, A-C) and three aptamer-peptide systems (1ETG, 1I9F, 1ULL, D-F).

EvoFlow-RNA-designed aptamers with the lowest RMSDs compared to wild-type aptamer analogs were selected for analysis. In each instance, designed aptamers bound to their small molecule (Figure 4 A-C) or peptide (Figure 4 D-F) targets. Unlike the similar binding interactions observed between designed and wild-type aptamers in the 7FHI and 1RAW systems, we observed a more significant binding disparity in the HIV small molecular inhibitor system (PDB ID: 2L94) despite the high structural homology between the aptamers (Fig 4B). We attributed this to sequence variations between the designed and wild-type aptamers that impacted pairwise binding interactions. We observed a similar phenomenon when analyzing aptamers designed against RSG peptide (Figure 4E). However, for HIV-rev peptide aptamers (Figure 4D,F), we observed that high structural homology to the wild-type aptamer resulted in more similar binding scores.

## Discussion

We introduced **EvoFlow-RNA**, a bidirectional RNA language model based on masked discrete diffusion that unifies representation learning and sequence generation. Trained on 30M ncRNA sequences, it achieves comparable performance with the leading BERT models, excelling in secondary structure prediction, and generating diverse, native-like RNA sequences. For RNA design, EvoFlow-RNA designs aptamer scaffolds while preserving functional motifs with sub-1Å RMSDs. Our results demonstrate the effectiveness of masked diffusion for RNA modeling, providing a scalable alternative to existing methods. Despite these results, we observed a few key limitations that could be items for future work. First, we observed that successful performance was not observed across all representation learning tasks, with EvoFlow-RNA performing below expectations on mRNA-related tasks. Though this was expected given that the model was trained entirely on ncRNA, its subpar performance on distance map prediction (DMP) is peculiar given its excellent ability on secondary structure and contact map prediction.

Second, though we observed that EvoFlow-RNA-generated libraries most closely mirrored native ncRNAs across metrics for sequence composition and biophysical properties, the pLDDT distributions did not always align closely. Often, EvoFlow-RNA generated sequences with pLDDT’s above those computed for native sequences. We note that this may be a consequence of the folding model. Lastly, we acknowledge that experimental validation for any generated RNAs, particularly the scaffolds, has yet to be done. The real-world applications using this model largely concern its generative capabilities, which can only be conclusively assessed in the lab. Our motif scaffolding experiments demonstrated a possible application of EvoFlow-RNA with a tractable validation method. If such results hold true experimentally, the possibility of ncRNA optimization can be seamlessly applied to the growing field of aptamer and tRNA therapeutics. We leave these items for future works.

## Code Availability

Our code is available at https://github.com/AtomBio/evoflow-rna.

## Author contributions statement

S.P. proposed the idea and conducted all the generation and benchmark experiments described in the paper. F.Z.P. outlined the mathematical framework and proposed the sampling algorithm experiments. K.F performed the molecular docking experiments. A.D.F. advised on nucleic acid biochemistry and intermolecular interactions. S.P., F.Z.P. and K.F. jointly wrote the manuscript, with all others contributing edits. P.C. and S.Y. supervised the project.

## Supplementary Figures

**Table A1.** EvoFlow-RNA model configurations.

| Hyperparameter | EvoFlow-RNA-33M | EvoFlow-RNA-150M | EvoFlow-RNA-650M |
| --- | --- | --- | --- |
| **Layers** | 12 | 30 | 33 |
| **Embed. dim.** | 480 | 640 | 1280 |
| **Attention heads** | 20 | 20 | 20 |
| **RoPE** | TRUE | TRUE | TRUE |
| **Attention dropout** | 0.1 | 0.1 | 0.1 |
| **Residual dropout** | 0.1 | 0.1 | 0.1 |

**Fig. A1.**
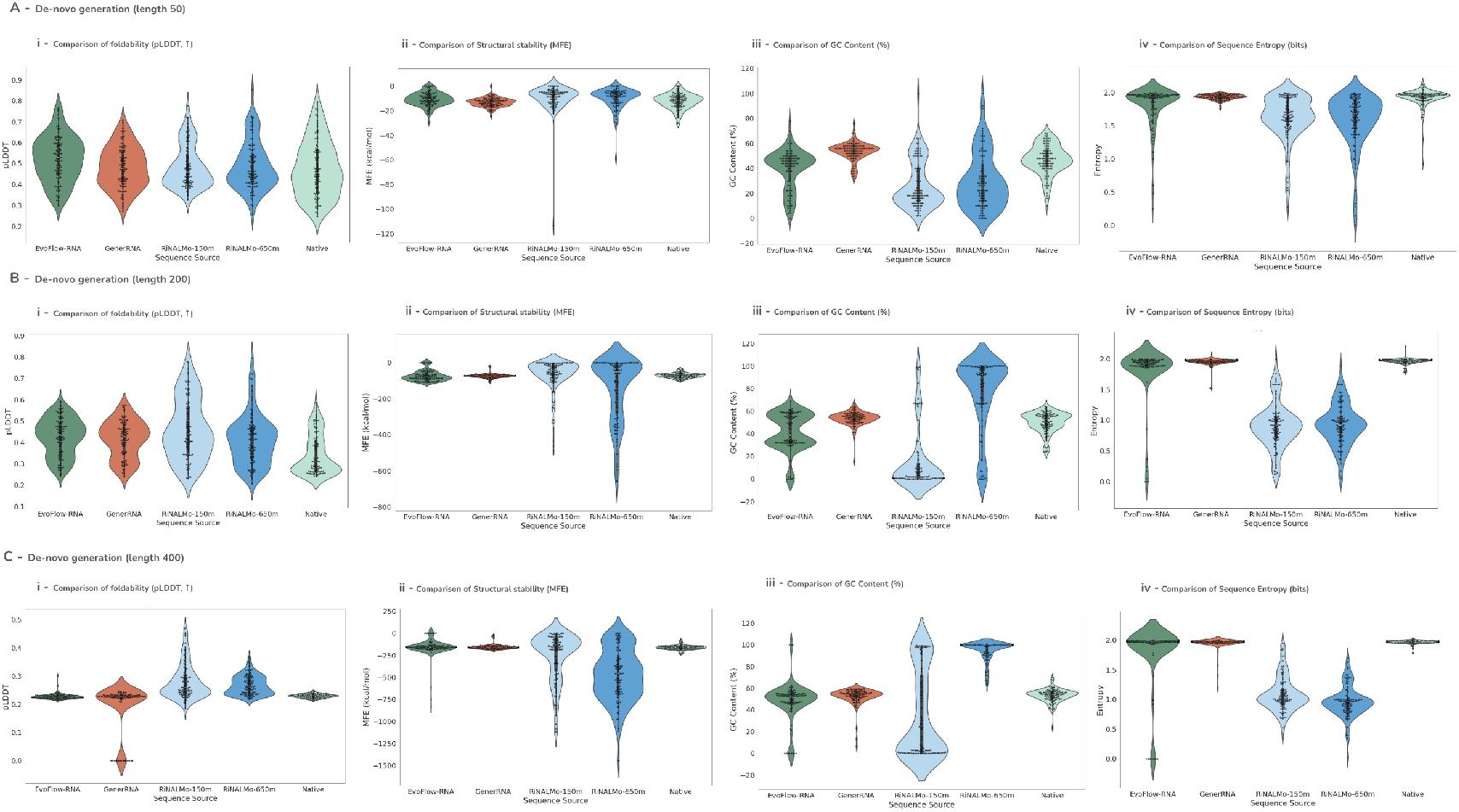
Unconditional generation evaluation with P2 (RDM, *η* = 1.0) for libraries of several lengths. Metric distributions are shown for libraries of length A) 50, B) 200, and C) 400.

**Table A2.** Ablation of EvoFlow-RNA model sizes and associated performance on bpRNA secondary structure prediction.

| Model | F-1 Score (↑) |
| --- | --- |
| EvoFlow-RNA-33M | 68.92 |
| EvoFlow-RNA-150M | 74.51 |
| EvoFlow-RNA-650M | 76.17 |

**Fig. A2.**
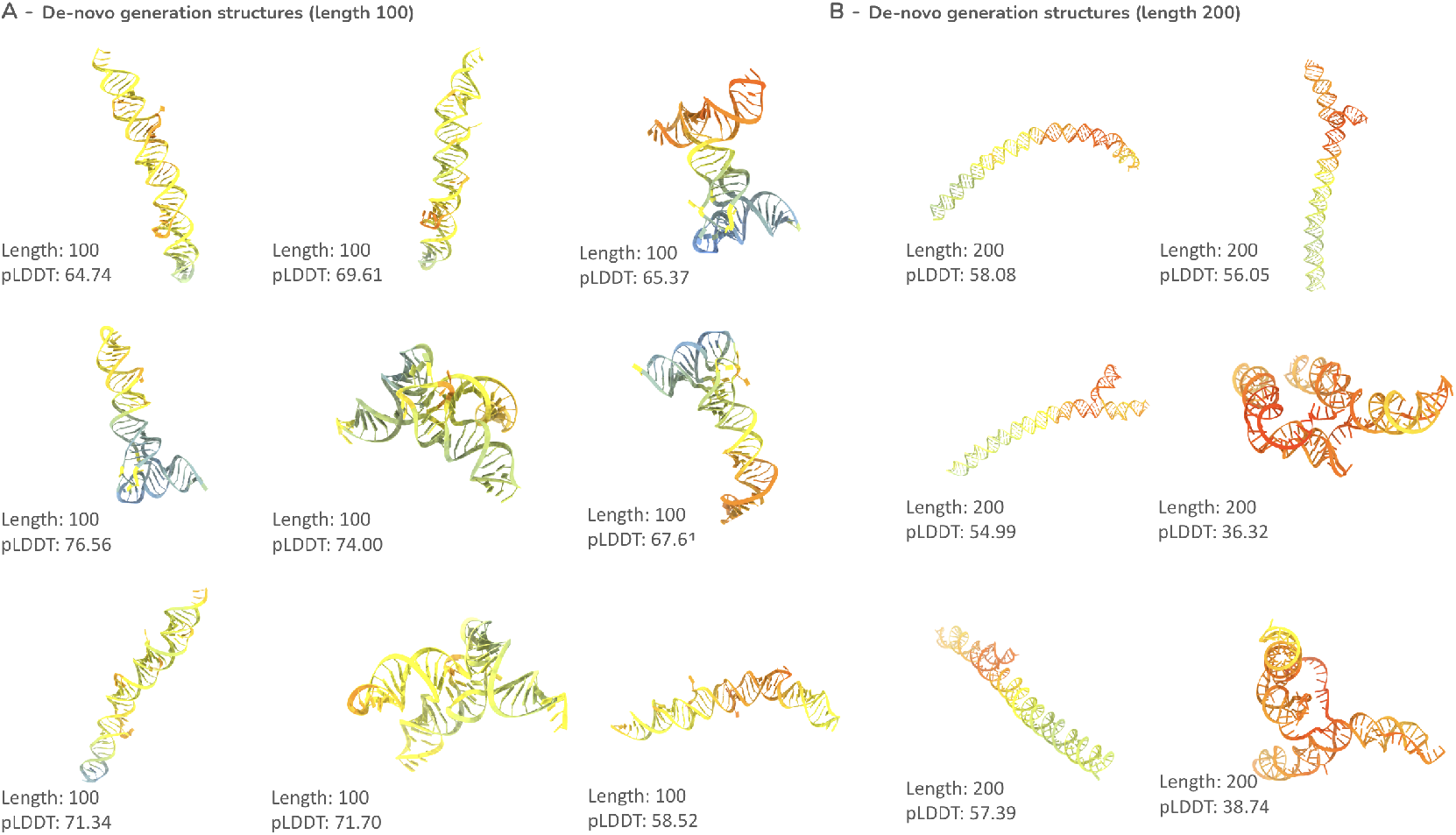
Additional structures for ncRNAs generated unconditionally by EvoFlow-RNA of length A) 100 and B) 200.

**Fig. A3.**
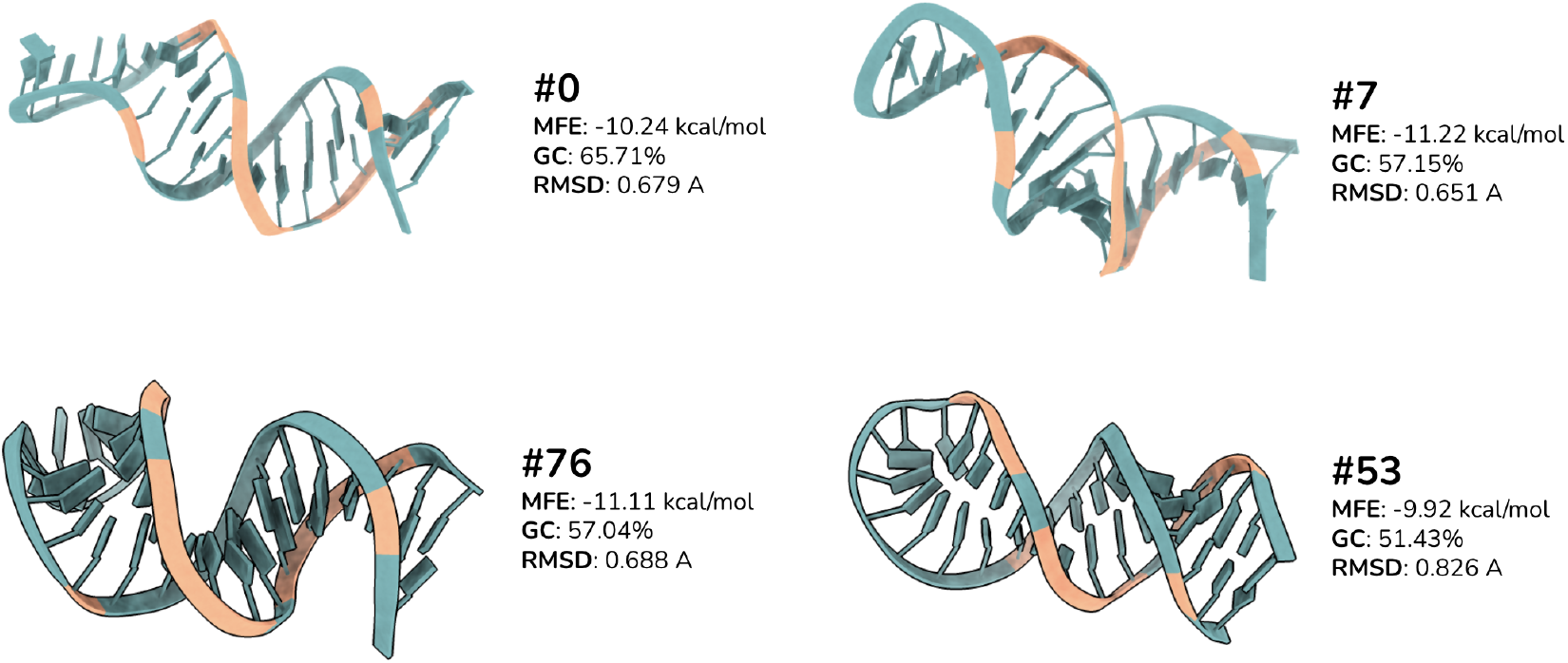
Additional scaffolds generated from the 1ULL HIV-1 Rev peptide binding aptamer.

